# Stem cell differentiation trajectories in *Hydra* resolved at single-cell resolution

**DOI:** 10.1101/460154

**Authors:** Stefan Siebert, Jeffrey A. Farrell, Jack F. Cazet, Yashodara L. Abeykoon, Abby S. Primack, Christine E. Schnitzler, Celina E. Juliano

## Abstract

The adult *Hydra* polyp continuously renews all of its cells using three separate stem cell populations, but the genetic pathways enabling homeostatic tissue maintenance are not well understood. We used Drop-seq to sequence transcriptomes of 24,985 single *Hydra* cells and identified the molecular signatures of a broad spectrum of cell states, from stem cells to terminally differentiated cells. We constructed differentiation trajectories for each cell lineage and identified the transcription factors expressed along these trajectories, thus creating a comprehensive molecular map of all developmental lineages in the adult animal. We unexpectedly found that neuron and gland cell differentiation transits through a common progenitor state, suggesting a shared evolutionary history for these secretory cell types. Finally, we have built the first gene expression map of the *Hydra* nervous system. By producing a comprehensive molecular description of the adult *Hydra* polyp, we have generated a resource for addressing fundamental questions regarding the evolution of developmental processes and nervous system function.

## Introduction

Hydrozoans have fascinated biologists for hundreds of years and have been at the center of fundamental discoveries in developmental biology, including animal regeneration (1) and stem cells. In fact, hydrozoan stem cells were the first to be described (2, 3). Among hydrozoans, stem cells and their differentiation pathways are best understood in the freshwater polyp *Hydra*, which have a relatively simple tissue structure and a small number of cell types. The stem cell populations, cell types, and lineage relationships are well characterized (4-9), however, the molecular characteristics of these cell types and the dynamics of their expression as they differentiate are largely unknown. We use single-cell RNA sequencing (scRNA-seq) to complement the extensive knowledge of *Hydra* developmental processes with unprecedented molecular characterization of fate specification pathways, thus revealing novel insights into the molecular programs driving cellular differentiation and patterning.

Homeostatic maintenance of the adult *Hydra* polyp results in the continual activity of every differentiation pathway, with all cells being replaced approximately every 20 days (10) (Fig. 1A-D). This requires the coordination of three distinct cell lineages (4-9) – endodermal epithelial, ectodermal epithelial, and interstitial. Each of these lineages is supported by its own stem cell population (Fig. 1A-D) (11). Ectodermal and endodermal epithelial cells in the body column are mitotic unipotent stem cells that self-replenish, resulting in continual displacement of these cells toward the extremities, where they are eventually shed (Fig. 1A) (12). Body column epithelial cells differentiate to build the basal disk at the aboral end or the hypostome and tentacles at the oral end (Fig. 1A,C). The interstitial cell lineage is supported by multipotent interstitial stem cells (ISCs) (13), which reside among the body column ectodermal epithelial cells (Fig. 1B). The adult *Hydra* has germline stem cells (GSCs) that self-maintain in a homeostatic animal, but ISCs can replace experimentally depleted GSCs (8, 14). ISCs give rise to three somatic cell types — nematocytes, neurons and gland cells. Nematocytes are single-use; neurons and gland cells are closely associated with epithelial cells and thus are continually displaced (7, 9). Despite the cell displacement and loss of interstitial lineage cells, cell numbers remain constant due to the following compensation mechanisms: 1) ISCs are mitotic and replace their own populations (15), 2) ISC progeny continually differentiate into the cell types of the interstitial lineage (15, 16), and 3) as differentiated neurons and gland cells are displaced towards the polyp’s extremities they change gene expression to reflect their location (17-19). Thus cell identity in *Hydra*depends on coordinating stem cell differentiation and gene expression programs in a manner dependent on the cell’s location along the oral-aboral body axis. In addition, for interstitial cells, identity also depends on the epithelium in which the cells reside. Understanding the molecular mechanisms underlying cellular differentiation and patterning in *Hydra* would be greatly facilitated by the creation of a spatial and temporal map of gene expression. Importantly, cnidarians hold an informative position on the phylogenetic tree as sister group to all bilaterians (20) and largely have the same complement of gene families found in vertebrates, thus offering the opportunity to identify conserved developmental mechanisms (21–23).

**Figure 1.**
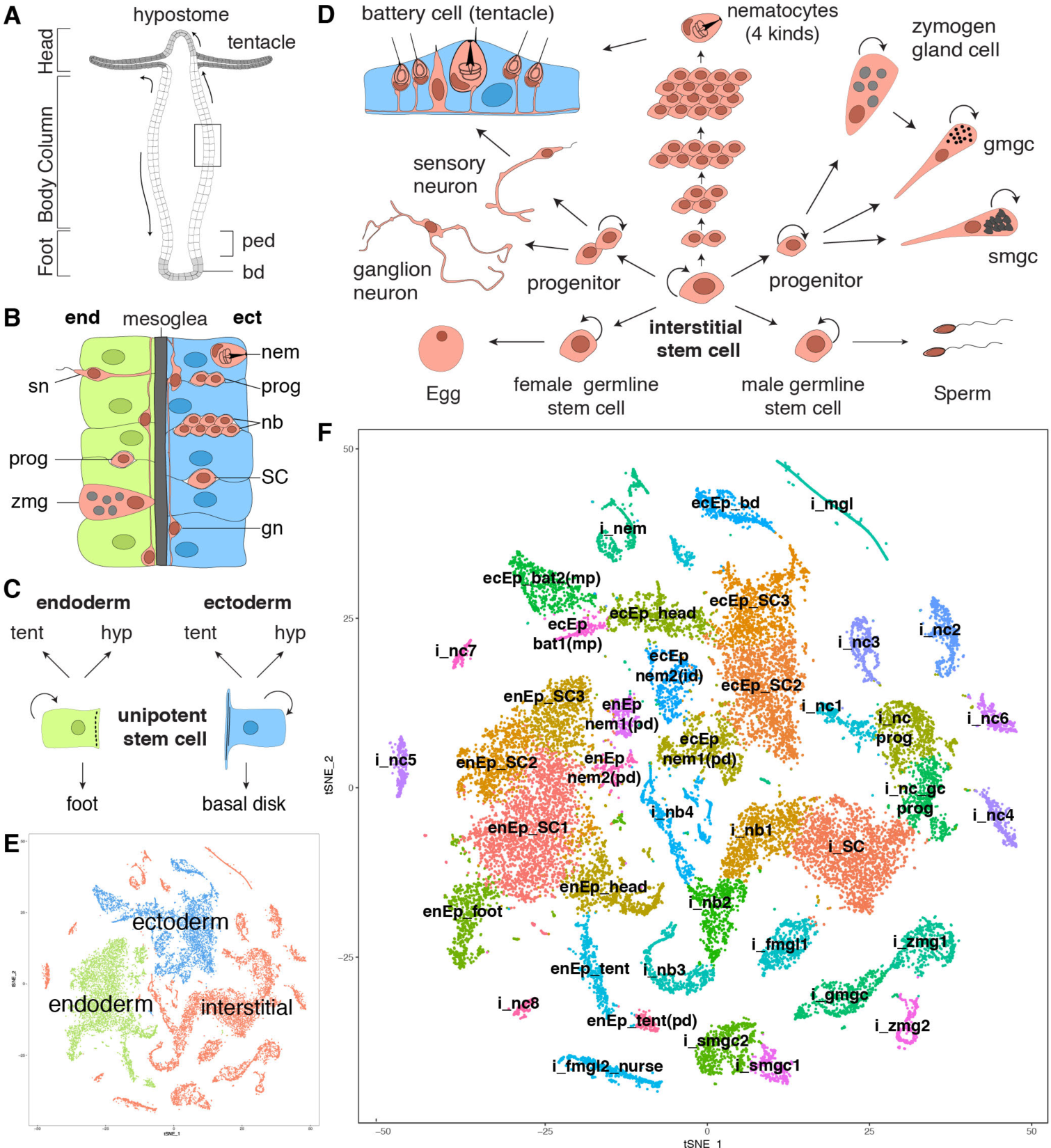
*Hydra* tissue composition and single cell RNA sequencing of 24,985 *Hydra* cells. A) The *Hydra* body is a hollow tube with a foot for attachment at the aboral end (bd: basal disc, ped: peduncle) and a head with a ring of tentacles (tent) at the oral end. The mouth opening arises at the tip of a cone shaped protrusion — the hypostome (hyp). B) Enlargement of box in A. The body column consists of two epithelial layers, the endoderm and the ectoderm, separated by an extracellular matrix — the mesoglea. Cells of the interstitial cell lineage reside in interstitial space between epithelial cells, except gland cells which are integrated into the endodermal epithelium. gn: ganglion neuron, SC: stem cell, sn: sensory neuron. C) Epithelial cells of the body column are mitotic, have stem cell properties, and give rise to terminally differentiated cells of the hypostome (hyp), tentacles (tent), and foot. D) Schematic of the interstitial stem cell lineage. The lineage is supported by a multipotent interstitial stem cell (ISC) that gives rise to neurons, gland cells, and nematocytes; ISCs are also capable of replenishing germline stem cells if they are lost. E) t-SNE representation of clustered cells colored by cell lineage. F) t-SNE representation of clustered cells annotated with cell state. Figures A-D adapted from (87). bat: battery cell, db: doublet, ecEP: ectodermal epithelial cell, enEP: endodermal epithelial cell, fmgl: female germ-line, gc: gland cell, gmgc: granular mucous gland cell, i: cell of the interstitial lineage, id: integration doublet, mgl: male germline, mp: multiplet, nb: nematoblast, nc: neuronal cell, nem: nematocyte, pd: suspected phagocytosis doublet, prog: progenitor, SC: stem cell, smgc: spumous mucous gland cell, tent: tentacle, zmg: zymogen gland cell.

In this study, we collect ~25,000 *Hydra* single-cell transcriptomes covering a wide range of cellular differentiation states. With these data, we built stem cell differentiation trajectories for each lineage and identified genes expressed in specific cell differentiation pathways, including the identification of putative regulatory modules that drive cell state specification. We identified a progenitor state common to the gland and neural differentiation pathways and explore gene expression of this cell state. We also revealed gene expression changes that occur as cells are displaced along the oral-aboral axis and as interstitial cells move into the endodermal layer. Finally, we have generated a molecular map of the nervous system with spatial resolution, which provides new opportunities to study mechanisms of neural network plasticity. We anticipate that providing a comprehensive molecular map as a resource to the developmental biology and neuroscience communities will rapidly advance our ability to make fundamental discoveries using *Hydra*.

## Results

### Single-cell RNA sequencing of whole Hydra reveals cell state transitions

Thirteen Drop-seq runs were performed on single cells from dissociated whole adult *Hydra* and two neuron enriched libraries were prepared using FACS to collect GFP-positive neurons from a transgenic line (Figs. S1,2 Tables S1,2). We mapped sequencing reads to a reference transcriptome and filtered for cells with 300-7,000 expressed genes and 500-50,000 Unique Molecular Identifiers (UMIs) resulting in a data set with a detected median of 1,936 genes and 5,672 UMIs per cell (Table S3). We clustered the cells and annotated cluster identity using published gene expression patterns (Figs. 1E,F, S3) and further validated identity by performing RNA in situ hybridization experiments (Fig. S4). In the clustering, cells separated according to cell lineage (Fig. 1E), and within each lineage, we observed the expected stem cell populations (Fig. 1F). We captured a large number of intermediate states between stem cells and differentiated cells and several differentiation trajectories are evident in the t-SNE representation, similar to what has been seen in scRNA-seq studies performed in planarians (24, 25). For example, clusters that correspond to differentiated head and foot epithelial cells are connected to their respective body column stem cell clusters (Fig. 1F). Additionally, the interstitial stem cell clusters are connected to both neuronal and nematocyte progenitors (nematoblasts). We also identified distinct clusters for differentiated cells of the interstitial lineage - neurons, gland cells, nematocytes, and germ cells (Fig. 1F). We applied non-negative matrix factorization (NMF) to the full dataset and subsequently to all lineage subsets to identify modules of genes that are co-expressed within cell populations (Fig. S5) (26); this approach has been used previously to identify co-expressed genes in scRNA-seq data sets (27). As described below, the recovered gene modules were used for doublet identification, trajectory characterization, and the identification of transcription factor binding sites enriched in the cis-regulatory elements of co-regulated genes.

Beside expected Drop-seq doublets (*i.e.* encapsulation of two cells in a single droplet), we identified additional doublet categories that are due to tight physical association between cells that provide novel insight into *Hydra* biology. First, battery cells, located in the tentacle ectoderm, consist of epithelial cells with integrated sensory neurons and nematocytes (28–31), and as expected, express epithelial, nematocyte, and neuronal markers (Figs. 1D,F “bat1” and “bat2”, S6). Second, some ectodermal cells along the body column house nematocytes (Fig. 1F ecEP nem2(id), Fig. S7) (28, 32). Third, neurons integrated in hypostomal epithelial cells have been reported (32). We found additional unexpected co-expression of interstitial and epithelial transcripts in single cells, including nematoblast gene expression in endodermal epithelial cells, an association that has not been reported (Fig. S7J-L). We attribute this, in part, to the ability of epithelial cells to phagocytose cells of the interstitial lineage, which occurs even across the extracellular matrix (mesoglea) that separates the ectodermal and endodermal epithelial layers (33–35). Indeed, we observed interstitial cells engulfed by epithelial cells in our dissociations using transgenic lines expressing fluorescent proteins (Fig. S2, S8). This supports our conclusion that prevalent phagocytosis generated many of the doublets in our data set. We addressed these instances of cell multiplets in downstream analyses and removed them from the data set prior to trajectory reconstruction.

### Trajectory reconstruction of epithelial cells reveals genes involved in axial patterning

To identify position-dependent gene expression patterns in epithelial cells along the body column, we performed trajectory analysis on subsets of endodermal and ectodermal epithelial cells using the R package URD (27) (Fig. 2A,B). URD connects cells with similar gene expression and uses simulated random walks to find gene expression trajectories between a terminal cell population and a starting progenitor cell population. This required that we remove doublets of all categories from the epithelial cell subsets, which we accomplished by implementing a novel approach using NMF co-expression modules to identify doublet signatures (see supplementary methods for details, Fig. S7). In vivo, epithelial cells divide in the body column and are displaced toward the extremities (Fig. 1A); however, to order endodermal and ectodermal cells according to expression state along the body column (the oral-aboral axis), we generated branching trajectories for each lineage by defining the foot cells (aboral) as the starting point (root) and hypostome and tentacle cells (oral) as two separate endpoints. To validate our endodermal and ectodermal epithelial differentiation trajectories, we visualized the expression of well-characterized genes (Fig. 2C,D, S9). For example, for the ectodermal trajectory we found that *HyAlx*, *Hym301*, *HvTSP*, *and Wnt9/10c* recapitulated published expression patterns in our data (Fig. 2C, S9A) (36–39) and we validated basal disk expression for a previously uncharacterized ectodermal gene t29450 (Fig. 2C, Fig. S4I). For the endodermal trajectory, we found that *CnNK-2*, *budhead*, *HyBra1*, and *Wnt3* recapitulated published expression patterns in our data, and we validated endodermal expression for previously uncharacterized genes *NAS14* (t13067) and *t2741* (Fig. 2D,E,F) (40–42).

**Figure 2.**
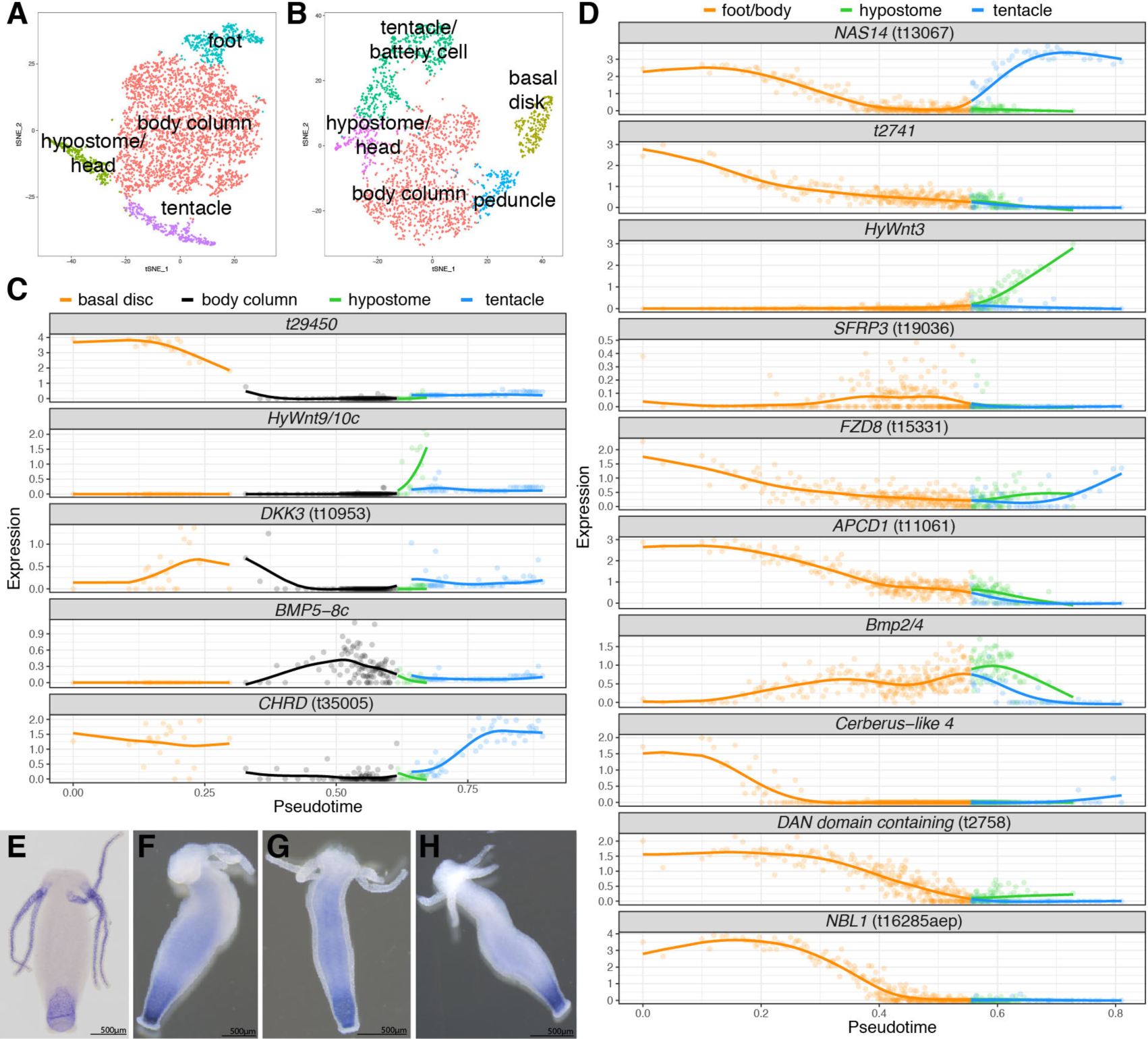
Differential gene expression along the oral-aboral axis reveals genes involved in axial patterning. A) t-SNE representation of subclustered endodermal epithelial cells and B) subclustered ectodermal epithelial cells. C) Trajectory plots of cells ordered by gene expression state along the body column (aboral to oral) for ectodermal (C) or endodermal epithelial cells (D). (C) Trajectory plots for genes expressed in ectodermal epithelial cells. *t29450* (Fig. S4I), *HyWnt9/10c*(39), *DKK3* (t10953), *BMP5-8c* (88), *CHRD* (t35005). (D) Trajectory plots for genes expressed in endodermal epithelial cells. *NAS14* (t13067), *t2741*, *HyWnt3*(41), *SFRP3* (t19036), *FZD8* (t15331), *APCD1* (t11061), *Bmp2/4*(88), *Cerberus-like 4* (88), DAN domain containing gene *t2758*, *NBL1* (t16285aep). E-H) Endodermal expression patterns obtained using RNA in situ hybridization consistent with predicted patterns. E) *NAS14* (t13067), F) *t2741*, G) *FZD8* (t15331), H) *APCD1* (t11061).

In cnidarians, Wnt genes are major players in axial patterning and integral parts of the oral organizer (39, 41, 43, 44). We identified multiple previously uncharacterized Wnt antagonists with highest expression in the *Hydra*foot. In the endodermal trajectory, two Frizzled related genes, *FZD8* (t15331) and *SFRP3* (t19036), and a *APCD1* homolog (t11061) have graded expression originating from the aboral end (Fig. 2D,G,H). In the ectodermal trajectory, we identified a previously uncharacterized *Dickkopf*gene, *DKK3*(t10953), with expression in the aboral epithelial cells and elevated expression in a subset of tentacle cells. The spatial expression of these four putative Wnt antagonists at the aboral end of the animal is consistent with a function in repressing Wnt signals that emanate from the oral organizer (Fig. 2C,D). We also interrogated our data set for genes involved in BMP signalling because the integration of Wnt and BMP signalling is a conserved aspect of patterning in vertebrates (45–47). We found several putative BMP antagonists with highest expression in the foot and often mutually exclusive expression domains with BMP ligands (Figs. 2C,D, S9A,B). In the ectoderm, an uncharacterized chordin-like gene CHRD (t35005) was identified with expression in the basal disk and tentacles that was strikingly complementary to BMP5-8c expression in cells of the body column. Therefore, this analysis suggests genes with candidate roles in integrating positional information during axis specification in cnidarians that merit future functional analysis. Finally, we used these trajectories to generate comprehensive sets of genes with variable expression along the oral-aboral axis in the epithelial lineages (Fig. S10). In addition, NMF analyses for cells of the epithelial subsets revealed gene modules differentially expressed along the body column (26) (Figs. S11,12). These spatially and temporally resolved gene expression profiles for epithelial cells of the body column that differentiate into oral and aboral structures are a valuable resource for determining regulators of epithelial cell terminal differentiation, such as transcription factors and signaling molecules.

### Identification of multipotent interstitial stem cells and trajectory reconstruction of the interstitial lineage

We extracted 12,470 interstitial cells from the whole data set (Fig. 1E), performed subclustering, and annotated the clusters (Fig. 3A, Fig. S13). The tSNE representation of subclustered interstitial cells clearly showed ISC differentiation into neurons and nematocytes (Fig. 3A). To build differentiation trajectories using URD required that we define a population of cells as the root (i.e. the multipotent ISCs). We identified markers specific to nematogenesis and neurogenesis; *HvSoxC* is expressed during both processes, suggesting this gene as a marker for interstitial differentiation (Fig. 3B, S13). Interestingly, we observed a conspicuous population of cells without *HvSoxC* expression (Fig. 3B), and we hypothesized that these *HvSoxC*-negative cells are the multipotent ISCs. We attempted to identify transcripts specific to the *HvSoxC*-negative population, but these cells are largely devoid of specific markers. Interestingly, this is consistent with planarian pluripotent cNeoblasts, which are similarly defined by an absence of cell-type specific genes (24). The forkhead transcription factor *FOXL1* (t12642) is expressed in the *HvSoxC*-negative population and continues to be expressed throughout nematogenesis, but its expression ceases during neurogenesis (Fig. 3C). Therefore, to build differentiation trajectories for cells of the interstitial lineage, we defined cells that are predominantly *HvSoxC-*negative/*FOXL1*-positive as the root (i.e. multipotent ISC population) (Fig. S14). We recovered a branching tree of interstitial stem cell differentiation that resolves neurogenesis, nematogenesis, and gland cell differentiation (Fig. 3D). We performed double fluorescent in situ hybridization (FISH) on several interstitial lineage genes to validate the trajectory tree (Fig. S15). Many of the gene modules previously identified by NMF analysis were specific to each differentiation pathway with ordered expression in pseudotime (Fig. S16), thus revealing with unprecedented detail the gene expression changes that underlie differentiation in the interstitial lineage.

**Figure 3.**
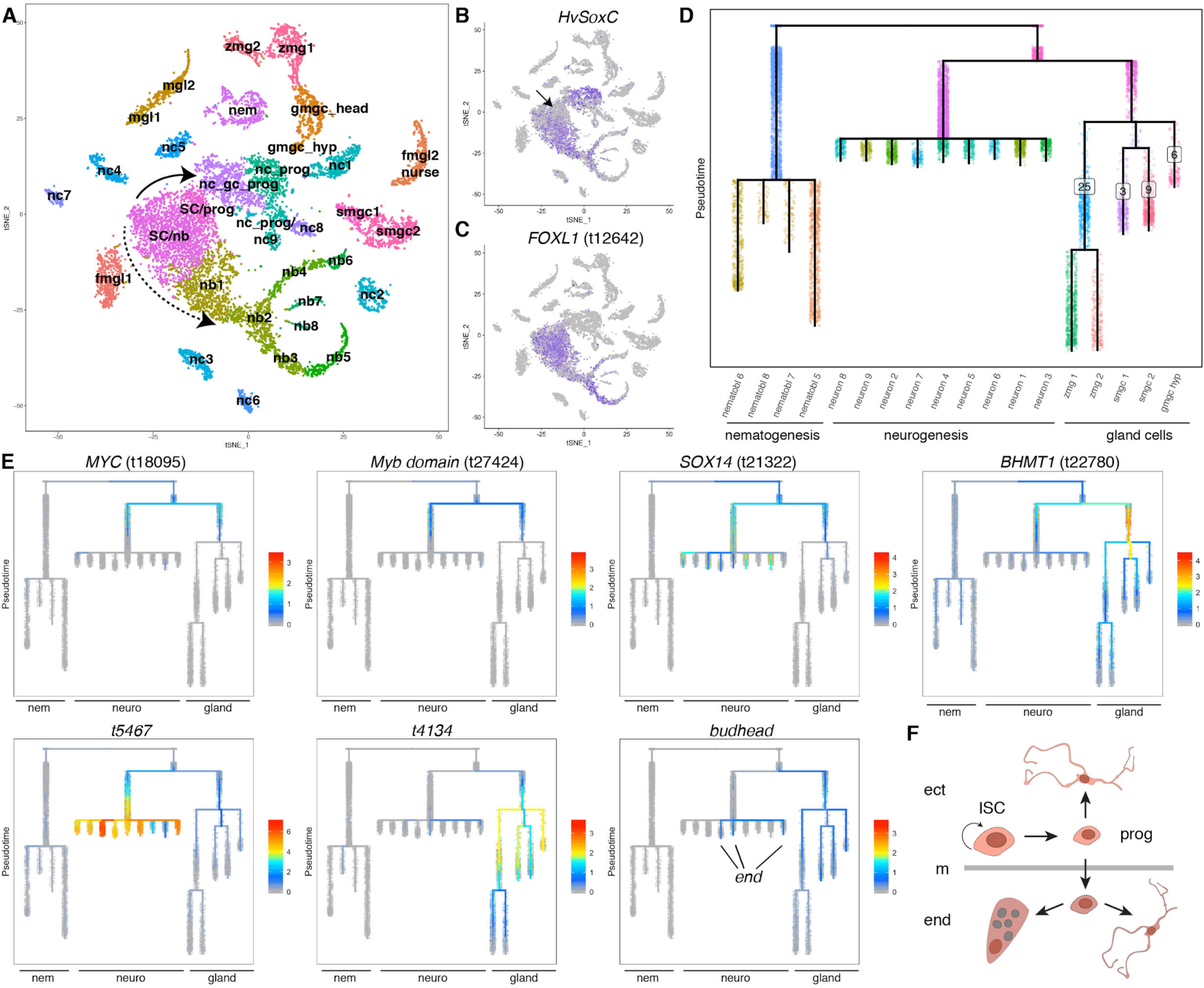
Trajectory analysis of the interstitial lineage suggests a bipotential neuron/gland cell progenitor. A) t-SNE representation of interstitial cells with clusters labeled by cell state. Solid arrow: neurogenesis/gland cell differentiation. Dashed arrow: nematogenesis. fmgl: female germ-line, gmgc: granular mucous cell, mgl: male germline, nc: neuronal cell, nb: nematoblast, nem: nematocyte, smgc: spumous mucous cell, prog: progenitor, SC: stem cell, zmg: zymogen gland cell. B) *HvSoxC* expressed in progenitor cells. Arrow indicates putative ISC population, which is negative for *HvSoxC* expression. C) Forkhead transcription *FOXL1*(t12642) factor expressed in ISCs and nematoblasts. D) URD differentiation tree of the interstitial lineage (see Fig. S14 for segment numbering). E) Gene expression in the shared progenitor state and during neurogenesis/gland cell differentiation. *MYC* (t18095) and Myb domain containing gene *t27424* are expressed in the shared progenitor state and in early neurogenesis/gland differentiation. *SOX14*(t21322) and *BHMT1* (t22780) have overlapping expression domains with *MYC*/*t27424* but neurogenesis or gland specific expression in later stages. Uncharacterized genes t5467 and t4134 with partially overlapping domains with *SOX14/BHMT1* but neurogenesis or gland specific expression. Forkhead transcription factor budhead is expressed in cells localized in the endoderm. end: endodermal neurons. F) Model for progenitor specification. Ectodermal ISCs give rise to a progenitor that can give rise to ectodermal neurons. Progenitors that translocate into the endoderm acquire endodermal gene expression and are able to give rise to glands or neurons. ect: ectoderm, end: endoderm, ISC: multipotent stem cell, m: mesoglea, prog: progenitor.

### Trajectory analysis of interstitial cells suggests a common neuron and gland cell progenitor

The trajectory analysis of the interstitial lineage revealed that the early nematogenesis differentiation program is distinct from early neurogenesis and gland cell formation. By contrast, neuron and gland differentiation transits through a previously undescribed shared progenitor cell state (Fig. 3D). We identified genes that are expressed in the common progenitor, including a *MYC* homolog (t18095) and a Myb domain containing gene *t27424* (Figs. 3E, S15). In addition, we find evidence that both neural and gland cell specific genes are simultaneously activated in the common progenitor. For example, *SOX14* (t21322) is activated in the common progenitor and is highly expressed during neurogenesis, but lost as cells differentiate into gland cells (Fig. 3E). Similarly, the gland cell gene *BHMT1* (t22780) is also activated in the common progenitor, but expression is lost during neurogenesis (Fig. 3E). We also found *MYC* (t18095) co-expressed in cells with expression of markers that are exclusive to each lineage such as *t5467* (neuron) or *t4134* (gland) (Fig. 3E). Finally, neuronal and gland-specific gene expression modules, recovered from our NMF analysis, are expressed in the common progenitor (Fig. S16 I,J). In addition, we identified a set of genes that are broadly expressed in endodermal epithelial cells and in cells from the interstitial lineage that are located in the endoderm. For example, the forkhead transcription factor *budhead*was originally described to be expressed in endodermal epithelial cells (48), but we also found expression in a subset of the neural/gland common progenitor cells, in cells of the endodermal gland cell trajectories, in a subset of neuron progenitors, and in differentiated endodermal neurons (Fig. 3E, see below for identification of endodermal neurons). Thus, *budhead* expression is found in all cell types that reside in the endodermal layer and we found several additional genes with correlated expression (Fig. S17). We therefore propose the existence of a dual primed neural/gland progenitor, which initially arises from the ISCs restricted to the ectoderm, and then acquires endoderm specific gene expression as the progenitor enters the endodermal epithelium and subsequently gives rise to neurons and gland cells (Fig. 3F). Future work should test the existence of this state and elucidate the molecular mechanism underlying fate decisions.

### Subtrajectory analyses of interstitial cell types

We next set out to explore specification and plasticity of different cell types within the interstitial lineage (Fig. 1D). First, we examined nematocytes, which contain one of the most complex eukaryote organelles - the nematocyst (49). These are used to sting and immobilize prey. *Hydra* nematocytes can each have one of four types of nematocysts: desmonemes, holotrichous and atrichous isorhizas, and stenoteles (28, 50). Forty-six percent of the cells in *Hydra* are nematoblasts (the cells that differentiate to become nematocytes) or nematocytes (51). These cells need constant replacement, so nematogenesis is the most prominent differentiation event in *Hydra*. We identified one cluster (Fig. 3A, “nem”) containing fully differentiated nematocytes based on expression of known mature nematocyte markers, but could not unambiguously assign four expected terminal fates, and therefore this cluster was excluded from the trajectory analysis. This ambiguity may be caused by similar transcriptional profiles across nematocyte types once the nematocyst has formed. However, in situ hybridizations for genes specifically expressed in the cluster positively identified two subpopulations – nematocytes that harbor differentiated stenoteles (Fig. 4A-C) and nematocytes that harbor differentiated desmonemes (Fig. 4D-F). Both the t-SNE representation (Fig. 3A) and URD analysis (Fig. 3D) revealed a prominent nematocyte trajectory with four branches that we hypothesize correspond to four nematoblast types, harboring developing nematocysts of the different kinds. We identified cells of two of these branches as differentiating desmonemes or stenoteles, based on co-expression of genes with the identified subpopulations in the differentiated nematocyte cluster (Fig. 4G,H). Our desmoneme assignment is further supported by absence of *nematogalectin A* expression, which is consistent with previous expression studies (52) (Fig. 4I). Expression of proliferating cell nuclear antigen (PCNA, t10355) suggested that cells within the four branches are post-mitotic (Fig. 4J). We identified transcription factors differentially expressed at the trajectory termini and performed in situ hybridization and found that these genes are expressed in nests of cells within the body column (S18). This demonstrated that our trajectories contain only nematoblasts because these cells undergo incomplete cytokinesis and only resolve into single nematocytes once they mature (53) (Fig. 1D). While there has been extensive work on nematocyst diversity, facilitated by their extreme morphological and functional differentiation, very little is known about nematocyte molecular diversity and how different the cells that harbor different types of nematocysts are. The identification of genes that are differentially expressed between nematocytes harboring different nematocyst types provides a toe-hold into understanding the specification and construction of these extraordinary organelles, which are the defining feature of Cnidaria.

**Figure 4.**
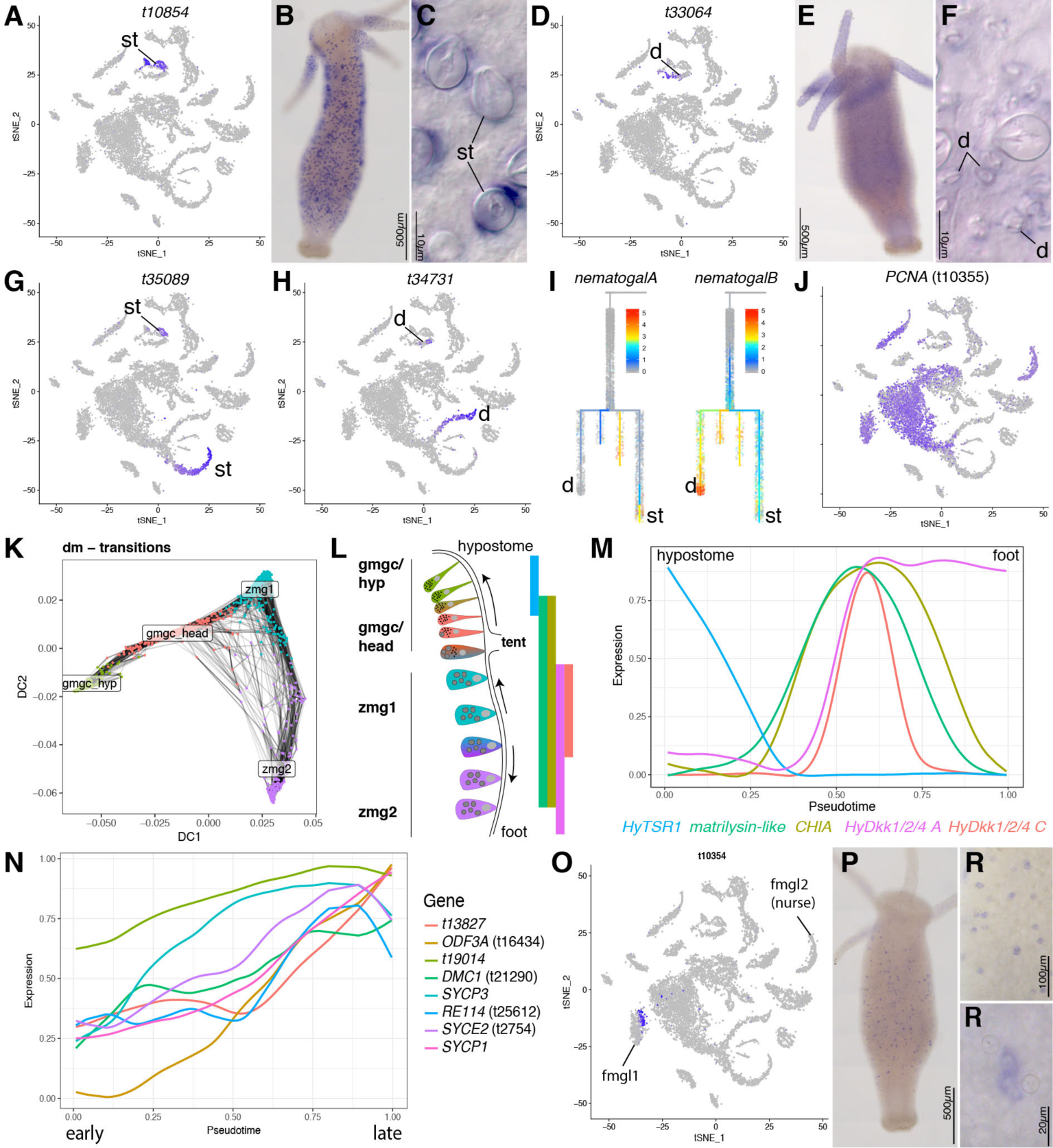
Subtrajectory analyses of interstitial cell types. A-I) Identification of trajectories for nematocytes forming stenoteles or desmonemes. A-F) Expression plots and RNA in situ hybridizations for transcripts *t10854*and *t33064*. A-C) *t10854* is expressed in stenoteles (st). D-F) *t33064* is expressed in desmonemes (d). G) *t35089* expression linking trajectory branch to stenotele fate. H) *t34731* expression linking trajectory branch to desmoneme fate. I) Lack of *nematogalectin A* expression supports desmoneme nematocyst type (52). *Nematogalectin B*is expressed in all trajectories. J) PCNA expression marks proliferative progenitor cells. K) Diffusion map for gMGC/ZMG gland cells. Transition probabilities (lines) suggest that zmg2 cells can differentiate directly into head gland cells (gmgc_head), but also more prominent transitions from zmg1 into zmg2 or zmg1 into gmgc of the head (linear trajectory). L) Model for linear ZMG/gMGC location dependent changes. Gland cells that are displaced change expression and morphology. Colors of cells correspond to populations depicted in (K). Bars show known expression domains for genes depicted in (M). tent: tentacle. M) URD linear ZMG/gMGC trajectory recapitulates known gene expression along the body column. N) URD reconstruction of the male germline shows meiosis genes peaking at the end of the trajectory. O-R) Putative female germline stem cell marker *Hvfem-2*(t10354). O) Plot showing expression in a subset of cells in the early female cluster. P-R) *Hvfem-2* is expressed in single cells or pairs scattered within the body column.

Second, we analyzed gland cells, which are interspersed between endodermal epithelial cells and are constituent parts of the endodermal epithelium. The two types of mucous gland cells (MGCs) predominantly found in the head are granular MGCs (gMGC) and spumous MGCs (sMGC) (Fig. 3A) (54, 55). Zymogen gland cells (ZMG) are found throughout the body and location-dependent changes in gland cells occur as they are displaced along the body column (56, 57). This includes ZMGs turning into gMGCs when they are displaced into the head {Siebert:2008kf}. Therefore, the maintenance of gland cell populations in *Hydra* occurs by both location-dependent changes and stem cell differentiation, which is reflected in our analyses. In the URD reconstruction of the interstitial lineage, we recovered differentiation of ISCs into all gland cell types found in the head (Fig. 3D, segments 25, 3, 9, 6, Fig. S19). While ISCs also differentiate into body column gland cells, we did not observe this; we hypothesize that this reflects a high rate of gland cell differentiation in the head, while body column gland cells perhaps primarily renew themselves mitotically. We recover the cell state continuum between ZMGs and gMGCs as a connection between the gMGCs (segment 25) and two separate populations of ZMGs (Fig. 3D). This trans-differentiation is represented in reverse in the reconstruction—*in vivo*, gMGCs would be the endpoint when body column ZMGs are displaced into the head (58). However, because we did not observe transitions between the ISCs and body column ZMGs, we instead used ZMGs as the endpoint for the reconstruction. Interestingly, the analysis suggested that there are two distinct ZMG cell states in the body column which is consistent with previous findings (56, 57). We next explored the transition probabilities between cell states directly (Fig. 4K). Our URD analysis suggested that both distinct ZMG states can change directly into a gMGC, but that it is more common for the cell state characteristic of the upper body column (zmg1) to do so. To capture the more dominant relationship, where body column ZMGs (zmg1) transition into either gMGCs or lower body column ZMGs (zmg2), we reconstructed a linear trajectory along the oral-aboral axis (Figs. 4L,M, S20). Known gland cell expression patterns along the body column (Fig. 4M) were recapitulated in this trajectory, and numerous new spatially varying genes in the population were revealed (Fig. S10). Analogously, we constructed a linear trajectory for the spumous gland cell populations in the *Hydra* head (Figs. S10, S21). Overall, our analysis reveals a broad range of gland cell states in *Hydra* that can be achieved through multiple molecular paths; future experiments should address the factors that control which molecular trajectory individual cells take and the downstream consequences for those cells.

Finally, we explored the germ cell clusters recovered in our data set. Although we used non-sexual polyps to build our Drop-seq libraries in most cases, four of the libraries included animals with either testes or developing eggs. We excluded germline cells from the interstitial lineage tree reconstruction because differentiation of GSCs from ISCs does not typically occur in a homeostatic animal, so we did not expect to observe transition states linking ISCs to GSCs. We, however, did elucidate the spermatogenesis trajectory by analyzing the progression of cell states found in the two male germline clusters that were recovered in the subclustering of interstitial cells (Figs. 4N, S10, S22). To evaluate the validity of this trajectory, we determined the expression patterns of genes with conserved function in meiosis and sperm development. The expression of synaptonemal complex proteins, *DMC1*(t21290aep), and sperm tail protein *ODF3A*(t16434aep) peak at the end of the trajectory (Fig. 4N, S22). The expression of meiotic genes at the end of the trajectory suggests that we captured spermatogonia and spermatocytes, but that spermatids were likely not captured due to low transcript abundance in mature sperm. Finally, we identified and confirmed several new genes expressed exclusively in germ cells, including male germline-specific histone proteins (Fig. S23).

We identified two female germ cell clusters, which likely correspond to early and late female germ cell development (Fig. 3A). Interestingly, all libraries, except the male-only library, contributed cells to the early female germline cluster, suggesting that early female germline cells are present in asexual polyps. By contrast, cells present in the late female germline cluster originated almost exclusively from libraries produced from animals with developing eggs. This late cluster is likely composed of nurse cells, since the majority of cells produced during oogenesis are nurse cells that are engulfed by the single oocyte (59). Cells in the late female germline cluster have the highest gene and UMI numbers compared to all other clusters in the dataset, which likely reflects deposition of maternal transcripts into the egg (Table S4). We performed in situ hybridizations for genes expressed in a subset of cells found in the early female germline cluster and found positive cells scattered as single cells and pairs of cells throughout the body column which may correspond to germline stem cells (Fig. 4O-R, S23). If so, this would be the first report of gene expression in *Hydra* that is specific to GSCs, as conserved germline genes (*vasa*, *nanos*, and *piwi*) are also expressed in ISCs (60–62). These data will therefore allow for the study of GSCs in *Hydra* through the construction of GSC reporter transgenic lines.

### Identification of regulatory modules that underlie cell type specification

The construction of lineage-specific differentiation trajectories allows us to determine the spatial and temporal expression patterns of transcription factors and thus gain insight into the gene regulatory networks that control cell type specification in *Hydra*. In our transcriptome assembly, we identified 435 transcripts with a DNA binding domain based on Pfam annotation, 424 of which are present in our scRNA-seq data set. As a step toward discovering the regulatory modules that control differentiation in *Hydra*, we aimed to identify the transcription factor binding sites shared by co-expressed genes and candidate transcription factors that could bind these sites. To accomplish this, we mapped our Drop-seq data set to the *Hydra*2.0 genome so that we could extract the regulatory sequences of co-expressed genes (63). Mapping *Hydra vulgaris* AEP sequencing reads to the *Hydra vulgaris* 105 genome, a closely related strain, rather than to our *Hydra vulgaris* AEP transcriptome, resulted in a reduction of the average number of mapped reads per library from 78.4% to 60.5%, but otherwise produced comparable results (Fig. S24).

To identify co-expressed genes, we used NMF to interrogate our genome-mapped data set and found 58 metagenes (*i.e.* sets of co-expressed genes), which are comparable to the set of metagenes identified in the transcriptome analysis (Fig. S25). To identify the putative regulatory regions of these co-expressed gene sets, we performed ATAC-seq on whole *Hydra vulgaris*105 polyps (64). We identified regions of locally enriched ATAC-seq read density (peaks) – signifying regions of open chromatin – and restricted our analysis to peaks within 5 kb upstream of the start codon of the genes in each NMF metagene. We then performed motif enrichment analysis to identify common transcription factor binding sites that may control the expression of genes belonging to a metagene. We found at least one significantly enriched motif for each of 39 metagenes. Notably, these metagenes had distinct sets of enriched motifs, suggesting potential differences in the transcription factor classes underlying various cell states (Fig. 5A, Fig. S26). For example, during nematogenesis, the paired box (Pax) motif is enriched in promoters of genes expressed during early and mid-stages, the forkhead (Fox) motif is enriched at mid- and late stages, and the POU motif is enriched only in late stages. The B-cell factor (EBF) motif is enriched in the female germline and the TCF motif is enriched in neurons and gland cells. Among epithelial cell states, motif enrichment is less tightly restricted to particular cell states. However, we did find that the ETS domain binding motif is enriched in metagenes expressed in the extremities (tentacles and foot) of both the endoderm and ectoderm. Additionally, homeodomain (Otx and Arx) and bZip motifs were enriched throughout both epithelial lineages and forkhead motifs appeared associated with genes expressed in endodermal epithelial cells (Fig. 5A). The enrichment of forkhead motifs in *Hydra* endoderm and *Nematostella* digestive filaments is consistent with a conserved function for forkhead transcription factors in cnidarian endodermal fate specification that is also found across bilaterians (65–67).

**Figure 5.**
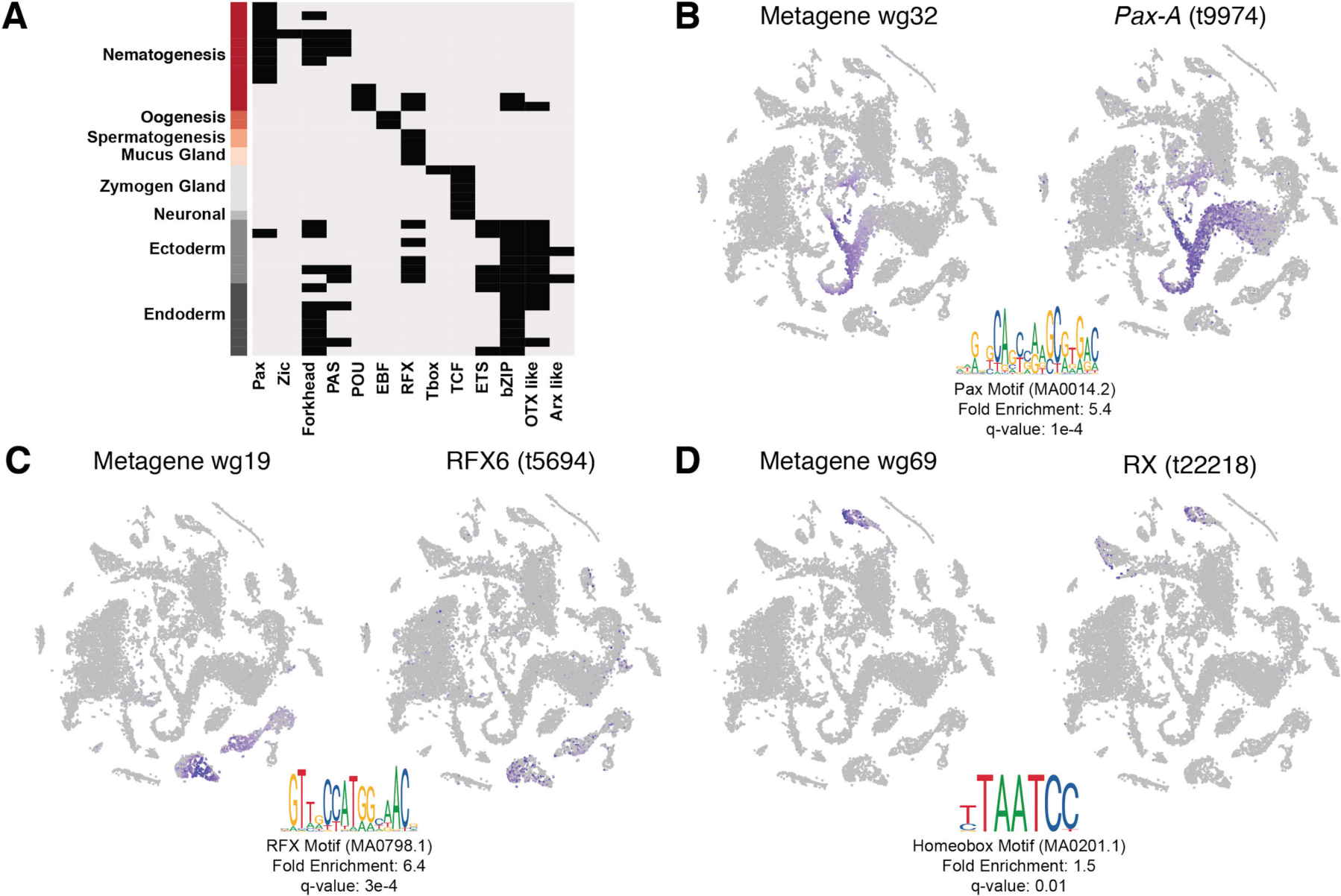
Motif enrichment analysis for gene modules and identification of candidate regulators. A) Enriched motifs (columns) found in open chromatin of 5’ cis-regulatory regions of co-expressed gene sets (metagenes) for listed cell states (rows). B-D) Metagene scores visualized on the t-SNE representation (left), significantly enriched motif found in 5’ cis-regulatory regions (bottom) and candidate regulators likely to bind identified motif with correlated expression (right). B) Metagene expressed during nematogenesis and putative PAX regulator. C) Metagene expressed in gland cells and putative RFX regulator. D) Metagene expressed in ectodermal epithelial cells of the foot and and putative homeobox regulator.

To determine the regulatory factors that may be coordinating these gene co-expression programs, we next identified transcription factors within each metagene that are predicted to interact with the binding site(s) enriched in that metagene using a combination of Pfam domain annotation and profile inference (JASPAR) (Fig. S27). For 24 of the 39 metagenes with enriched binding motifs, we found one or multiple candidate transcription factors with putative function in cell fate specification (Table S5). For example, we find a metagene (wg27) that consists of 144 genes co-expressed during nematogenesis. A Pax transcription factor binding motif was significantly enriched in potential regulatory regions of open chromatin near those genes, and the *Pax-A* transcription factor (t9974) is part of the metagene (Fig. 5A,B, Fig. S26). Our results therefore strongly suggest that *Pax-A* functions during early nematogenesis. This is concordant with a recent finding that *Pax-A* is required for nematogenesis during *Nematostella* development (65, 68). Similarly, we also found evidence that suggests that an RX homeobox transcription factor (t22218) functions in basal disk development and an RFX transcription factor (t5694) functions in gland cell specification; the latter was also reported for *Nematostella* (Fig. 5C,D) (65). For cases where we find more than one TF that is both expressed in the proper context and predicted to bind an enriched motif—such as the basal disk—we provide all transcription factors that met our criteria as candidate regulators (Table S5). Overall, we identified several candidates for key regulators of *Hydra* cell fate specification.

### A molecular map of the Hydra nervous system

The *Hydra* nervous system consists of two nerve nets, one embedded in the ectodermal epithelial layer and one embedded in the endodermal epithelial layer. Neurons are concentrated at the oral and aboral ends of the polyp (51). In a homeostatic animal, the neurons are displaced along the oral-aboral axis together with the epithelial cells; thus, neurons are lost and the nervous system must be continually rebuilt. Maintenance of the nervous system requires both the creation of new neurons from differentiating ISCs and changes in neuronal gene expression as neurons change location (17–19). Many molecular markers for neurons in the ectodermal nerve net have been identified, including neuropeptides and transcription factors (69–73), but no molecular markers for neurons in the endodermal nerve net have been described. We used our single cell data to identify the complete set of molecularly distinct neuronal subtypes in *Hydra* and determined their in situ location.

To determine the neuronal subtypes present in *Hydra*, we extracted neural progenitors and differentiated neurons from the dataset for subcluster analysis. We identified 15 clusters: three clusters consist of neuronal progenitor cells (Fig. 6A, clusters 0, 11, and 13) and the remaining 12 clusters are differentiated neuronal subtypes. To place these 12 neuronal subtypes into the ectodermal or endodermal nerve net, we performed TagSeq (74) on separated tissue layers that contained epithelium-associated neurons and conducted differential gene expression (DGE) analysis to identify genes with significantly higher expression in the endodermal or ectodermal epithelial layer (Fig. S28). Since the neurons remained attached to the epithelia, differentially expressed genes between the ectodermal and endodermal samples included neuron-specific genes, and the presence of these genes allowed us to score our neuronal clusters as ectodermal or endodermal. We clearly identified three endodermal neuronal clusters (2, 3, 8) and nine ectodermal neuronal clusters (1,4-7,9,10,12,14) (Fig. 6A, Fig. S28).

**Figure 6.**
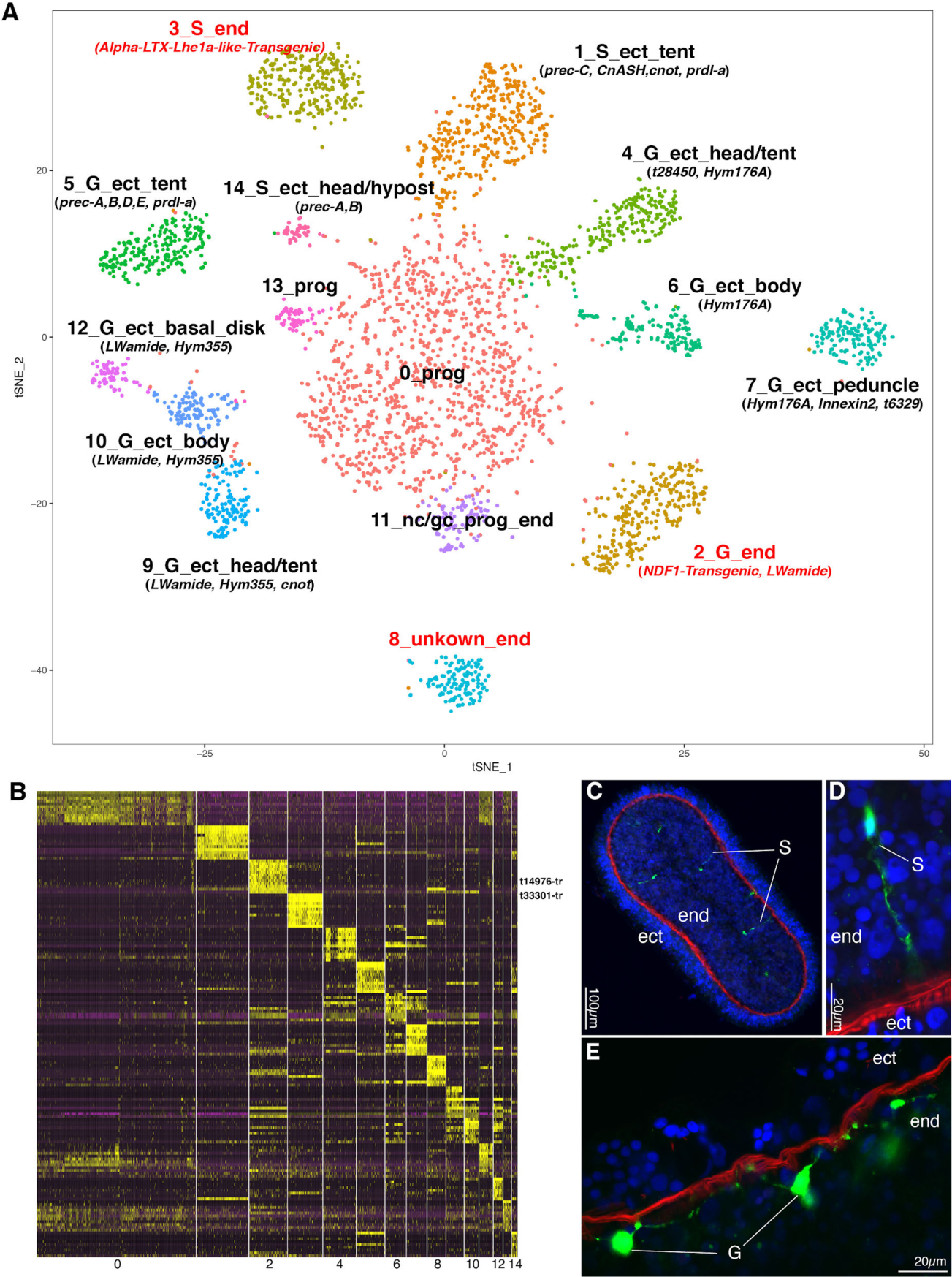
Molecular map of the *Hydra* nervous system with spatial resolution. A) Subclustering of neurons and neuronal progenitors. Cell states are annotated with cluster id, tentative neuronal subtype category — sensory (S) or ganglion (G), in vivo localization and gene markers used in annotations. B) Heatmap shows top twelve neuronal specific subtype markers. C-E) First molecular markers for endodermal neurons. C-D) Transgenic line (Alpha-LTX-Lhe1a-like(t33301)::GFP) expressing GFP in putative sensory neurons (cluster 3). E) Transgenic line (NDF1(t14976)::GFP) expressing GFP in endodermal ganglion neurons along the body column (cluster 2). Phalloidin staining (red) marks ECM and Hoechst (blue) marks nuclei. end: endoderm, ect: ectoderm, gc: gland cell, nc: neuronal cell, prog: progenitor.

To determine the location of the ectodermal neuronal subtypes along the oral-aboral axis, we used a combination of published in situ patterns, in situ patterns of our newly identified neuronal markers, and spatial information retained by the integration or phagocytic uptake of neuronal cells (see supplement for details) (Figs. 6A, S29,S30,S31). We predicted biomarkers for all neuronal subtypes (Figs. 6B, S32). To test the endodermal identity of clusters 2 and 3, we examined *NDF1* (t14976, specific to cluster 2) and*Alpha-LTX-Lhe1a-like* (t33301aep, specific to cluster 3) expression using GFP reporter lines. In case of NDF1, we found GFP expression in endodermal ganglion neurons in the entire body except tentacles (Fig. 6E). In the case of *Alpha-LTX-Lhe1a-like*, we found GFP expression in sensory neurons along the body column in the endoderm (Fig. 6C,D). Therefore our transgenic lines confirm endodermal localization of clusters 2 and 3 and demonstrate our ability to identify specific biomarkers for each neuronal subtype. In summary, we have produced a molecular map of the *Hydra* nervous system that, at the chosen resolution, describes 12 molecularly distinct neuronal subtypes and their in situ locations.

## Discussion

We present an extensively validated molecular map of *Hydra* cell states generated by sequencing approximately 25,000 single-cell transcriptomes collected largely from whole animals and supplemented with two neuron-enriched libraries. In addition, we provide differentiation trajectories for most of the cell populations in the animal. This gives us access to a catalog of transcription factors expressed at key developmental decision points and provides the first multi-lineage trajectory map of an adult animal. Several recent studies have demonstrated the value of conducting whole animal (24, 25, 65) or whole embryo scRNA-seq (27, 65, 75, 76) to uncover cell type diversity and the regulatory programs that drive cell type specification. Conducting scRNA-seq on a diversity of organisms will provide insights into the core regulatory modules underling cell type specification and the evolution of cellular diversity (77). Thus, our *Hydra* data set provides an additional opportunity for comparisons to be made in an evolutionary context.

Analysis of *Hydra* by single-cell RNA sequencing uncovered new technical challenges, and we provide new solutions that will likely be applicable to many systems. For example, *Hydra* epithelial cells are highly phagocytic (33, 78), which may be an important homeostatic maintenance mechanism to remove unwanted cells (33). Phagocytic cell uptake has been observed in variety of systems, and thus will likely present a challenge for interpretation of scRNA-seq results in future studies (79–81). We implemented an approach that has been incorporated into URD, in which we use NMF as an unbiased method to identify anomalies in the data that likely represent cell doublets or phagocytic events. Additionally, we demonstrate that our cell doublets also provide a unique strength by identifying cells that are spatially proximal to each other. For instance, we were able to use spatially localized epithelial genes in combination with epithelial–neuron doublets to predict the in situ location of neuronal subtypes. We envision that our approaches could be applied to other systems and will be particularly useful in animals where existing expression data are limiting.

We present the first molecular map of a dynamic and regenerative nervous system, which opens the door to understanding the molecular basis of neuronal plasticity and regeneration. Of the twelve distinct neuronal subtypes we have identified, three (the endodermal neurons) were previously completely uncharacterized molecularly. Our molecular map was validated using previously known gene expression patterns and through the identification of new neuronal biomarkers. We further validated our identification of the elusive endodermal neurons using transgenic approaches. The results of the validation tests consistently agreed with the predictions from the single-cell data. Thus scRNA-seq is a highly robust technique for identifying cell type, cell location, and developmental trajectory in *Hydra*. Neuron subtype-specific transgenes will provide powerful tools for experimental perturbations to test neuronal function and nervous system regeneration. Three distinct neuronal circuits have been described in *Hydra*; two in the ectoderm — rhythmic potential 1 (RP1) and contraction burst (CB) — and one in the endoderm — rhythmic potential 2 (RP2) (82). These circuits are likely composed of ganglion neurons connected throughout the body. Based on the in situ locations of the ganglion neurons we identified, we propose the following: 1) the ganglion endodermal neurons that comprise cluster 2 (Fig. 6A,B) make up the RP2 circuit, 2) the *LWamide*- and *Hym-355*-positive ganglion neurons of clusters 9,11, and 12 make up the RP1 circuit, and 3) the *Hym-176*-positive ganglion neurons of clusters 4,6, and 7 make up the ectodermal CB circuit. This is supported by the observation that the RP1 circuit is active in the basal disc (cluster 12), whereas the CB circuit extends aborally only to the peduncle (cluster 7) (82). We have identified genes with expression specific to neuron subtypes and circuits. One intriguing example are seven innexin genes, which are involved in building gap junctions, with differential expression in the nervous system (83) (Fig S33,34). In addition, the availability of markers specific to neuronal subtypes will enable precise alterations to neural circuits. Nervous system function in such engineered animals can be tested using newly developed microfluidic tools that allow for simultaneous electrical and optical recordings in behaving animals (84).

The interstitial lineage differentiation trajectories presented here advance multiple questions that have long been central to the field. For example, we unexpectedly identified a cell state that is shared in the neuron and gland cell trajectory (Fig. 3D). It remains to be elucidated if this cell state is bipotential or rather the common progenitor pool is a collection of molecularly similar unipotent cells already determined to be either glands or neurons. Regardless, our data suggests a model in which multipotent ISCs first decide between a nematocyte or gland/neuron fate and then a second decision is made by the common gland/neuronal progenitor. This contrasts with previous models that posit a common progenitor with the capability of giving rise to neurons or nematocytes (85). First, we hypothesize that this shared developmental history between gland cells and neurons could point to a shared evolutionary history for these two secretory cells types. Second, these data suggest a model in which a bipotential gland/neuron progenitor born in the ectodermal layer, where multipotent ISCs reside, traverses the extracellular matrix to provide the endodermal layer with both gland cells and neurons (Fig. 3F); these are the only two interstitial cell types that populate the endodermal layer. Interestingly, we find that the forkhead transcription factor *budhead* (48) is expressed in all endodermal cell types – endodermal neurons, gland cells, and epithelial cells (Fig. S35). We therefore hypothesize that *budhead* is a determinative factor for endodermal location and future work will focus on testing this model through lineage tracing and gene function tests.

In summary, the construction of developmental trajectories at single cell resolution reveals the complete set of genes underlying cell fate decisions and thus advances major questions in developmental biology (27, 75, 76, 86). Adult *Hydra* polyps, which are in a constant state of development, enables the capture of all states of cellular differentiation using scRNA-seq. An important future goal is to use scRNA-seq to rapidly assess the effect of mutations on all cell types (27, 76, 86). *Hydra* has a diversity of fate specifications from multiple stem cell types, yet is simple enough to be completely captured by a relatively small number of sequenced single cells, which presents an exciting opportunity to begin such phenotyping efforts. The transcription factors that we identified at key developmental decision points are the most exciting first candidates to profile after perturbation. In conclusion, this resource and the experimental approaches we describe open doors in multiple fields including developmental biology, evolutionary biology, and neurobiology.

## Acknowledgements

We thank Robert Monroy for collection of separated endoderm and ectoderm epithelial layers for TagSeq and Ram Abhineet, Jamie Ho, Qianyan Li, and Noemi Sierra for wet lab validations by RNA in situ hybridization. We thank Rob Steele and Catherine Dana for kindly providing the eGreen (EF1-alpha promoter::GFP) transgenic line, Toshitaka Fujisawa, Xiaoming Zhang, and Yukihiko Noro for kindly providing the neuro:GFP transgenic line, and Thomas C. G. Bosch for kindly providing *Hydra vulgaris* AEP. We thank Lutz Froenicke, Vanessa Rashbrook, Siranoosh Ashtari, Emily Kumimoto of the UC Davis DNA Technologies Core for their expert assistance with library preparations and sequencing. The Olympus FV1000 confocal used in this study was purchased using NIH Shared Instrumentation Grant 1S10RR019266-01. We thank the MCB Light Microscopy Imaging Facility, which is a UC Davis Campus Core Research Facility, for the use of this microscope. Many thanks to Charlie David for providing numerous and plentiful insights on *Hydra* biology that helped with the interpretation of the data. We thank Bryan Teefy for helpful discussion and moral support. We thank the following people for critical reading of the manuscript: Alex Schier, Rob Steele, Gary Wessel, Bruce Draper, Stefan Materna, Jacob Robinson, and Casey Dunn.

## Author Contributions

S.S., J.A.F., J.F.C., and C.E.J. conceived the study; S.S., J.A.F., J.F.C., and C.E.J. wrote the paper with revisions by A.S.P, Y.A., C.E.S.; S.S. and J.A.F. collected single cell transcriptomes; Y.A., J.F.C., A.S.P., and S.S. performed ISH validation experiments; A.S.P. performed transgenic validation experiments of neurons and FACS sorting; J.F.C. performed ATAC-seq; S.S., J.A.F., and J.F.C. processed the raw data and conducted data analysis; S.S. and C.E.S. built transcriptome and gene models; J.A.F. built differentiation trajectories; J.F.C. and S.S. performed regulatory module analysis.

